# HLA-EpiCheck : A B-cell epitope prediction tool for HLA proteins using molecular dynamics simulation data

**DOI:** 10.1101/2023.12.18.572133

**Authors:** Diego Amaya-Ramirez, Magali Devriese, Romain Lhotte, Cedric Usureau, Malika Smail-Tabbone, Jean-Luc Taupin, Marie-Dominique Devignes

## Abstract

The Human Leukocyte Antigen (HLA) system is the main cause of organ transplant loss through the recognition of HLA proteins by Donor-Specific Antibodies (DSA). Therefore, the identification of potentially immunogenic epitopes is a key task to refine organ allocation and then to improve the survival of transplanted organs. Here, we present HLA-EpiCheck, a machine learning predictor for B-cell epitopes on HLA proteins that leverages an unprecedented dataset of high-quality molecular dynamics simulations of 207 HLA proteins. Candidate epitopes are represented as surface patches centered on solvent-accessible residues and described by a set of 18 descriptors. The descriptors include both static and dynamic properties, such as hydrophobicity, electrostatic charges, relative solvent-accessible surface area and side-chain flexibility. The HLA-EpiCheck was trained using an Extra Trees ensemble learning method and was compared to DiscoTope-3.0, a state-of-the-art B-cell epitope predictor. HLA-EpiCheck largely outperformed DiscoTope-3.0 in the task of predicting HLA epitopes. HLA-EpiCheck was also used to assess the epitope status of a subset of non-confirmed eplets. The predictions were compared to experimental data and a notable consistency was found. These results suggest that HLA-EpiCheck could be used to better define HLA matching between donor and recipient to reduce *de novo* DSA formation and graft rejection.

## 2 Introduction

The Human Leukocyte Antigen (HLA) system plays a key role in the activation of the human immune response. It is responsible for presenting peptides to T-lymphocytes from antigen processing in cells. However, this same system is mainly responsible for the loss of organ transplants. This is due, firstly, to the fact that these membrane proteins are expressed in almost all nucleated cells of the organism, i.e. they are practically omnipresent [1]. Secondly, the genes encoding these proteins are the most polymorphic genes in human. HLA genes are localised on chromosome 6, position 6p21.31 and are classified as class I for the A, B and C loci, and class II for the DR, DQ and DP loci. In addition, class II loci are more complex as they associate two genes that are both polymorphic for DQ (DQA1 and DQB1) and DP (DPA1 and DPB1). As a consequence, it is extremely difficult to find HLA identical donors and recipients outside siblings, and therefore transplantations are almost always performed across HLA polymorphisms that may later on trigger an immune response (both humoral and cellular) against the donor. In this paper we investigate the molecular mechanisms of humoral response, in particular the process of recognition of HLA proteins by antibodies, also known as donor-specific antibodies (DSA) for those that target the donor HLA antigens. For this purpose, we try to characterize B-cell epitopes on HLA proteins. B-cell epitopes are protein regions recognized and bound by B-cell produced antibodies. Understanding the characteristics of HLA B-cell epitopes and predicting their localization on HLA antigens is of key importance for improving the prevention of humoral response in transplantation. In this regard, and despite the numerous studies that have addressed the subject [2–9], we found so far no studies that have analyzed the HLA B-cell epitopes from a dynamic point of view, in particular through molecular dynamics (MD) simulations.

The initial interest in MD simulations relies on their ability to refine the 3D structures of proteins, especially to correct stereochemical errors. This technique has proven its usefulness by being integrated into the protocols for predicting the 3D structure of proteins [10–14] although it has also been used in the refinement of structures obtained experimentally by X-ray diffraction [15, 16]. Secondly, MD simulations allow to study the dynamic properties of proteins such as side-chain flexibility and solvent accessibility. A recent study suggests that side-chain flexibility is a key element in antigen-antibody recognition [17]. Additionally, MD simulations give access to conformations that are closer to those that occur under ambient conditions. Indeed, the structures obtained by X-ray diffraction present the protein in a crystallized conformation, which is not necessarily exactly the same under ambient conditions. This also applies to the structures predicted by tools such as AlphaFold v2 [18], as they have all been trained using structures from the Protein Data Bank (PDB) [19], which mostly consist of X-ray diffraction-resolved structures.

DSA are the main cause of graft loss in organ transplantation [20, 21]. Polymorphic residues on mismatched HLA molecules (between donor graft and recipient), known as eplets, are considered the key components of the epitopes recognized by DSA [22] (see Figure 1). Putative eplets are typically identified by analysis in human serum (e.g. patients enrolled in transplantation programs, pre and post transplant) of reactivity patterns of Luminex Single Antigen (LSA) bead assays. A LSA assay represents a multiplex assay capable of detecting antibody binding to close to 200 among the most frequent HLA proteins in the population, on a Luminex apparatus. Each HLA protein in the assay differs from the others by at least one amino acid polymorphism. The HLA Eplet Registry [23] is a database that gathers data on putative and antibody-verified eplets reported in the scientific literature. Antibody-verified eplets are those for which an experimental validation (e.g. with patients’ sera) has been performed, in contrast to the putative eplets (also known as non-confirmed eplets) which have mainly be deduced from alignments of HLA proteins sequences, and therefore have not been confirmed yet. Antibody-verified eplet mismatches have been demonstrated to correlate with DSA formation and graft survival [24, 25]. So predicting B-cell immunogenic epitopes in HLA proteins could lead to a better definition of HLA matching between donor and recipient to reduce *de novo* DSA formation and graft rejection.

**Figure 1:**
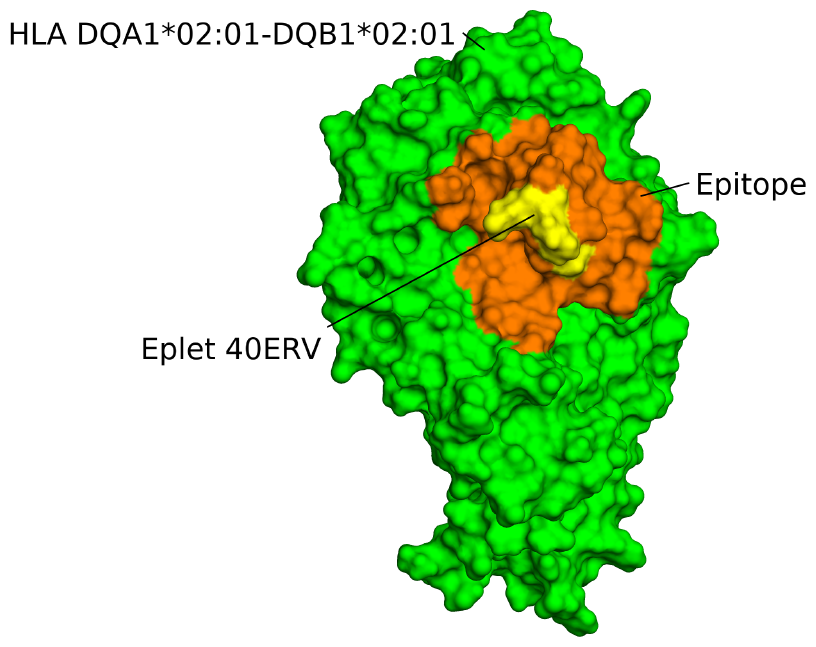
Visualisation of the eplet 40ERV and its associated epitope on the antigen HLA-DQA1*02:01-DQB1*02:01. Surface representation of antigen HLA-DQA1*02:01-DQB1*02:01. Eplet 40ERV (composed of three residues: 40E, 41R and 45V) is shown in yellow. The orange zone (patch) corresponds to residues that are 15Å at most away from residue 41R of the eplet 40ERV. This zone represents a putative epitope.

Thus, in the present work, we implemented a pipeline to generate an unprecedented dataset of high-quality MD simulations of 207 HLA antigens, hence covering all the antigens that are typically evaluated in LSA assays used in clinical practice. This dataset allowed us to train a machine learning (ML) predictor for B-cell epitopes on HLA proteins which we call HLA-EpiCheck. We leveraged the antibody-verified eplet database in the HLA Eplet Registry to train HLA-EpiCheck. Each candidate epitope is represented as a surface patch centered on a solvent-accessible residue and described by both static and dynamic types of descriptors.

We report here the performance of the HLA-EpiCheck predictor and compare it to a state-of-the-art B-cell epitope predictor (DiscoTope-3.0) [26]. Additionally, as several studies have demonstrated the major role of HLA-DQ in the development of *de novo* DSA in graft rejection [27–29], we used HLA-EpiCheck to assess the epitope status of a subset of non-confirmed eplets at the locus DQ and compared these results with experimental data.

Finally, we discuss the possible usage of HLA-EpiCheck in organ allocation to better characterize and rank donor-recipient pairs.

## 3 Results

Here, we first present the performance of our HLA-EpiCheck epitope predictor and an estimation of feature importance during the prediction task. Then, we evaluate the performance of our tool with respect to DiscoTope-3.0 [26]. Finally, we present a comparison with experimental results of a subset of non-confirmed eplets of HLA-DQ molecules.

### 3.1 Performance and feature importance of the HLA-EpiCheck predictor

#### 3.1.1 HLA-EpiCheck performance

Ten repetitions of 10-fold cross-validations were performed on the training set to evaluate the predictor performance. Precision, Recall and F1 metrics were used to evaluate performance for each class label (epitope, non-epitope), whereas AUC-ROC, AUC-PR and MCC metrics were used for global evaluation. The results in Tables 1, S5 and S6, and Figures S1 and S2 indicated a robust performance of the predictor as well as a good ability to discriminate the two labels (epitope/non-epitope). The average precision and recall over the 10 repetitions were 0.92 ± 0.025 and 0.837 ± 0.028 respectively for the epitope label and 0.934 ± 0.013 and 0.967 ± 0.01 respectively for the non-epitope label yielding average F1 score of 0.876 ± 0.019 and 0.95 ± 0.008 for the two labels respectively. The average AUC-ROC, AUC-PR and MCC were 0.979 ± 0.005, 0.958 ± 0.01 and 0.829 ± 0.025 respectively. Low standard deviation values indicate an homogeneous and robust performance of the predictor over the entire training set. For the epitope label, a better performance is observed on precision than on recall metric. This suggests that the predictor is better at predicting the epitope labels correctly than it is at identifying all of the epitope cases (precision is the proportion of predicted positive/epitope cases that are actually positive, while recall is the proportion of all positive/epitope cases that are correctly predicted). Therefore, a higher precision metric indicates that the tool is less likely to make false positives. On the other hand, the metrics AUC-ROC, AUC-PR and MCC were used to globally evaluate the performance of the predictor on the training set. A high performance was observed in the AUC-ROC and AUC-PR. This indicates that HLA-EpiCheck is performing well at distinguishing between the positive/epitope and negative/non-epitope labels. Similar to what was observed in the precision, recall and F1 metrics, the low values in the standard deviations on AUC-ROC and AUC-PR metrics suggest that our tool is robust. Regarding the MCC metric, although the values were lower than those obtained in the previously mentioned metrics, it is important to remember that this metric is less ≪ optimistic ≫ when evaluating a predictor because it takes into account the 4 components of the confusion matrix (i.e. true positives, true negatives, false positives, false negatives). Thus, the MCC result (0.829) denotes a very good performance of the tool. However, as suggested by the results on the recall metric, the predictor has room for improvement on false negatives.

**Table 1:** Performance evaluation of HLA-EpiCheck on the training set through 10 repetitions of 10-fold cross-validations.

| Metric | Epitope | Non-epitope |
| --- | --- | --- |
| | mean ± std | mean ± std |
| **Precision** | 0,92 ± 0,025 | 0,934 ± 0,013 |
| **Recall** | 0,837 ± 0,028 | 0,967 ± 0,010 |
| **F1** | 0,876 ± 0,019 | 0,950 ± 0,008 |
| **AUC-ROC** | 0,979 ± 0,005 | |
| **AUC-PR** | 0,958 ± 0,010 | |
| **MCC** | 0,829 ± 0,025 | |

#### 3.1.2 Feature importance

To evaluate the contribution of dynamic and static descriptors to the performance of HLA-EpiCheck, we calculated the Mean Decrease in Impurity (MDI) of each descriptor (see Figure 2). Simply put, the MDI is a normalized value that indicates the importance of a descriptor in the predictor performance. We found that the static descriptor H patch max is the most important contributor to performance (∼0.115, bottom bar in Figure 2). However, in second place is the dynamic descriptor F patch avg with a MDI of (∼0.082 which represents the average side-chain flexibility of the patch. In addition, the descriptors H central (hydrophobicity of the central residue) and H patch avg (average hydrophobicity of the patch) rank fifth and sixth respectively in the feature importance analysis (∼0.072 and ∼0.066 MDI values respectively). These results highlight the importance of hydrophobicity in the predictive ability of the tool (the sum of the MDI values of H patch max, H central and H patch avg is 0.253). Although 4 of the 5 descriptors with the highest MDI values are static descriptors, the sum of the MDI values of this type of descriptor is 0.47, indicating that although the dynamic descriptors do not stand out for their high MDI values, these descriptors have a very consistent mid-range contribution. However, the descriptor F patch avg stands out among the dynamic descriptors by having the second-highest MDI value. This suggests, as mentioned in [17], that side-chain flexibility plays a key role in antigen-antibody recognition.

**Figure 2:**
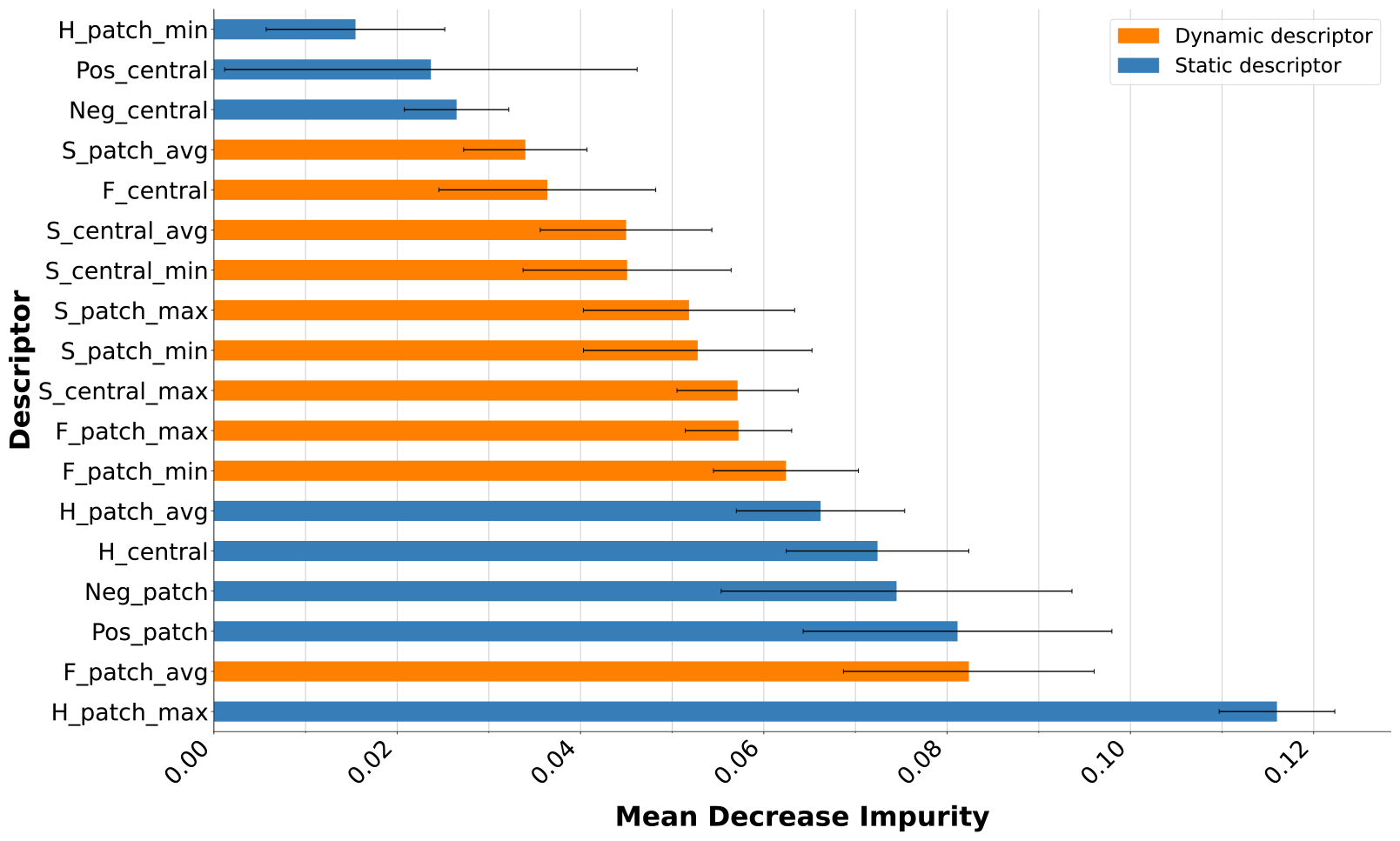
MDI feature importance using F1 metric. Horizontal bar length corresponds to mean MDI values (each tree in the forest having a MDI value). Orange bars correspond to dynamic descriptors. Blue bars correspond to static descriptors. Whiskers correspond to standard deviations of MDI values over the 100 trees.

### 3.2 Performance evaluation on the test set and comparison with DiscoTope-3.0

For the evaluation of our predictor, we first ran HLA-EpiCheck on our test set (i. e. unseen samples for HLA-EpiCheck) and we obtained the values presented in Table 2 (column HLA-EpiCheck). Concerning the results when considering all the loci, the precision and recall were 0.93 and 0.82 respectively for the epitope label, and 0.92 and 0.97 respectively for the non-epitope label yielding F1 scores of 0.87 and 0.95 for the two labels respectively. The AUC-ROC, AUC-PR and MCC were 0.98, 0.98 and 0.82 respectively. These results were similar to those obtained in the performance evaluation on the training set which indicates a robust and non overfitting predictor. We also found as in the training set evaluation, a better performance on precision metric than in recall metric for the epitope label which indicates again that the tool is less likely to make false positive predictions and has room for improvement on false negative predictions. Concerning the results per locus, HLA-EpiCheck showed a high performance for loci A, B, C and DP with F1 scores not lower than 0.86 for epitope prediction and 0.93 for non-epitope prediction. DQ popped up as the less-performing locus, especially for the non-epitope label with a F1 score of 0.65 while the epitope label remained high with a F1 score of 0.92. This can easily be explained by the small number of samples for the non-epitope label (only 95 samples, the lowest among all the loci). Interestingly, loci C and DR performed well despite being the loci with the lowest number of samples (counting both labels) with F1 scores of 0,9 and 0.79 respectively for the epitope label and, 0.95 and 0.74 respectively for the non-epitope label.

**Table 2:** Performance evaluation of HLA-EpiCheck and DiscoTope-3.0 on test set.

| Locus | Metric | HLA-EpiCheck |  | DiscoTope-3.0 |  |
| --- | --- | --- | --- | --- | --- |
|  |  | Epitope | Non-epitope | Epitope | Non-epitope |
| **A** | **Precision** | 0.89 | 0.91 | 0.31 | 0.63 |
|  | **Recall** | 0.83 | 0.95 | 0.32 | 0.62 |
|  | **F1** | 0.86 | 0.93 | 0.32 | 0.62 |
| **B** | **Precision** | 0.91 | 0.97 | 0.19 | 0.85 |
|  | **Recall** | 0.82 | 0.99 | 0.26 | 0.79 |
|  | **F1** | 0.87 | 0.98 | 0.22 | 0.82 |
| **C** | **Precision** | 1 | 0.91 | 0.5 | 0.75 |
|  | **Recall** | 0.81 | 1 | 0.58 | 0.68 |
|  | **F1** | 0.9 | 0.95 | 0.54 | 0.71 |
| **DP** | **Precision** | 0.97 | 0.94 | 0.32 | 0.72 |
|  | **Recall** | 0.85 | 0.99 | 0.46 | 0.59 |
|  | **F1** | 0.91 | 0.96 | 0.38 | 0.65 |
| **DQ** | **Precision** | 1 | 0.48 | 0.91 | 0.15 |
|  | **Recall** | 0.86 | 1 | 0.56 | 0.58 |
|  | **F1** | 0.92 | 0.65 | 0.69 | 0.24 |
| **DR** | **Precision** | 0.83 | 0.7 | 0.66 | 0.44 |
|  | **Recall** | 0.76 | 0.79 | 0.29 | 0.79 |
|  | **F1** | 0.79 | 0.74 | 0.4 | 0.56 |
| **All loci** | **Precision** | 0.93 | 0.92 | 0.56 | 0.8 |
|  | **Recall** | 0.82 | 0.97 | 0.55 | 0.8 |
|  | **F1-score** | 0.87 | 0.95 | 0.56 | 0.8 |
|  | **AUC-PR** | 0.98 |  | 0.67 |  |
|  | **AUC-ROC** | 0.98 |  | 0.76 |  |
|  | **MCC** | 0.82 |  | 0.35 |  |

Then, we compared HLA-EpiCheck against DiscoTope-3.0 (a state-of-the-art B-cell epitope prediction tool). Like our tool, DiscoTope-3.0 is a binary classifier that predicts whether a residue is part of an epitope or not. However, DiscoTope-3.0 does not consider the surrounding residues forming the 3D patch and is only trained with descriptors extracted from static 3D structures. The predictions of DiscoTope-3.0 were made on structures corresponding to the first frame of MD simulations. Although HLA-EpiCheck classifies patches rather than single residues, for comparison purposes, we will impute the predicted label of the patch to its central residue. The comparison was performed on the test set. Table 2 summarizes the performances of both tools. HLA-EpiCheck largely outperforms DiscoTope-3.0 with an MCC value of 0.83 for HLA-EpiCheck and only 0.35 for DiscoTope-3.0, especially in the prediction of the epitope label since the F1 score is 0.87 for HLA-EpiCheck and 0.56 for DiscoTope-3.0. The difference is not so strong for the non-epitope label which yields a F1 score of 0.95 for HLA-EpiCheck compared to 0.8 for DiscoTope-3.0. The Precision-Recall and ROC curves are shown in Figures S3 and S4. Concerning the results per locus, DiscoTope-3.0 performed acceptably in locus C with F1 scores of 0.54 and 0.71 for epitope and non-epitope predictions, respectively. However, the tool performed poorly in predicting the epitope label in loci A, B, DP and DR, with F1 score not greater than 0.4, while the ability to predict the non-epitope label was acceptable with F1 scores ranging from 0.56 (locus DR) to 0.82 (locus B). In the case of locus DQ, acceptable performance was observed for predicting the epitope label (F1 score of 0.69) while poor performance was observed for the non-epitope label (F1 score of 0.24). Interestingly, We observed the same pattern as in HLA-EpiCheck concerning the locus DQ, i.e., a significant drop in performance for the non-epitope label.

### 3.3 Comparison with experimental results on a subset of non-confirmed eplets

The HLA Eplet Registry contained 492 eplets (accessed February, 2023), of which only 146 had the antibody-verified status. We found that of the 56 non-confirmed eplets at the locus DQ in the HLA Eplet Registry (see Table S7), 40 eplets had at least one solvent-accessible residue. We computed the HLA-EpiCheck scores *𝒮*_*e*_ and *𝒮*_*e_*_*resid* for each of these 40 eplets (see Table 3). Remember that *𝒮*_*e*_ is an average over all HLA antigens presenting a given eplet of the averages of epitope predictions for each solvent-accessible residue composing this eplet. In other words, for a given eplet and a given antigen, an epitope prediction (1 for epitope and 0 for non-epitope) is given by HLA-EpiCheck to each patch centered on a solvent-accessible residue that is a member of the considered eplet on a given antigen. We first computed for each considered antigen the average of the epitope predictions over all the solvent-accessible residues of the eplet, and then the average of these eplet-level prediction scores over all antigens displaying this eplet.

**Table 3:** HLA-EpiCheck scores for the non-confirmed eplets of the locus DQ. Scores are also computed for single eplet residues. Max distance column corresponds to the maximum distance between eplet residues. Mean center of mass along the trajectory of residues is used for computing distances. Allele families DQA1*02, DQA1*04, DQA1*05 and DQA1*06 have a deletion relative to the remaining DQA1*01 and DQA1*03 allele families at position 56, so the residue numbering changes for these alleles. Eplets concerned with this deletion are marked with a *. Values for the HLA-EpiCheck score per eplet residue and Max distance are not shown for eplets composed of only one solvent-accessible residue. In 𝒮_*e_resid*_ column, missing residues of composite eplets are not solvent-accessible. Non-confirmed eplets experimentally validated are highlighted in the lightgreen rows.

| No | Eplet name | $\mathcal{S}_e$ | $\mathcal{S}_{e\_resid}$ | Chain | Max distance [Å] |
| --- | --- | --- | --- | --- | --- |
| 1 | 75IL | 1 | - | $\alpha$ | - |
| 2 | 67VT | 1 | - | $\beta$ | - |
| 3 | 56PS | 1 | - | $\beta$ | - |
| 4 | 56PD | 1 | 56P:1; 57D:1 | $\beta$ | 6.3 |
| 5 | 56PA | 1 | 56P:1; 57A:1 | $\beta$ | 5.5 |
| 6 | 56P | 1 | - | $\beta$ | - |
| 7 | 55RPD | 1 | 55R:1; 56P:1; 57D:1 | $\beta$ | 6.5 |
| 8 | 55PPD | 1 | 55P:1; 56P:1 | $\beta$ | 6.4 |
| 9 | 55PPA | 1 | 55P:1; 56P:1 | $\beta$ | 5.8 |
| 10 | 3P | 1 | - | $\beta$ | - |
| 11 | 185I | 1 | - | $\beta$ | - |
| 12 | 70GT | 1 | 67V:1; 70G:1 | $\beta$ | 7.3 |
| 13 | 70RT | 1 | - | $\beta$ | - |
| 14 | 135G | 1 | - | $\beta$ | - |
| 15 | 23R | 0.96 | - | $\beta$ | - |
| 16 | 66DR | 0.93 | 66D:0.86; 67I:1; 70R:1 | $\beta$ | 8.2 |
| 17 | 74EL | 0.91 | 74E:0.81; 77T:1 | $\beta$ | 8.6 |
| 18 | 66D | 0.86 | 66D:0.82; 67I:1 | $\beta$ | 5.2 |
| 19 | 67VG | 0.83 | 66E:0.5; 67V:1; 70G:1 | $\beta$ | 8.2 |
| 20 | 40ERV | 0.82 | 40E:0.86; 41R:0.71; 45V:0.92 | $\alpha$ | 8.3 |
| 21 | 185T | 0.80 | - | $\beta$ | - |
| 22 | 66ER | 0.79 | 66E:0.38; 67V:1; 70R:1 | $\beta$ | 6.8 |
| 23 | 66EV | 0.71 | 66E:0.42; 67V:1 | $\beta$ | 7.3 |
| 24 | 75I* | 0.67 | 74I:1; 75I:1; 160D:0.3; 161D:0.44; 162I:0.6; 163I:0.63 | $\alpha$ | 49.9 |
| 25 | 135D | 0.52 | - | $\beta$ | - |
| 26 | 23L | 0.50 | - | $\beta$ | - |
| 27 | 130R | 0.36 | - | $\beta$ | - |
| 28 | 167H | 0.33 | - | $\beta$ | - |
| 29 | 160AD* | 0.33 | 159A:0.2; 160A:0.31; 160D:0.3; 161D:0.46 | $\alpha$ | 5.3 |
| 30 | 160A* | 0.27 | 159A:0.23; 160A:0.31 | $\alpha$ | - |
| 31 | 3S | 0.17 | - | $\beta$ | - |
| 32 | 129QS | 0.14 | 129Q:0.14; 130S:0.14 | $\alpha$ | 6.3 |
| 33 | 167R | 0.13 | - | $\beta$ | - |
| 34 | 129H* | 0.04 | 128H:0.07; 129H:0 | $\alpha$ | - |
| 35 | 175E* | 0 | 174E:0; 175E:0 | $\alpha$ | - |
| 36 | 160S | 0 | - | $\alpha$ | - |
| 37 | 160D | 0 | - | $\alpha$ | - |
| 38 | 130Q | 0 | - | $\beta$ | - |
| 39 | 130A | 0 | - | $\alpha$ | - |
| 40 | 125G | 0 | - | $\beta$ | - |

In Table 3, 14 of the 40 eplets have a score of 1, 7 eplets have a score in [0.8, 1[, 3 eplets have a score in [0.6, 0.8[, 2 eplets have a score in [0.4, 0.6[, 4 eplets have a score in [0.2, 0.4[, 4 eplets have a score in] 0, 0.2[] and 6 eplets have a score of 0. The HLA-EpiCheck scores have been compared with experimental results obtained for 15 non-confirmed eplets in order to establish their epitope status (antibody-verified). These experimental results consist of adsorption of patients’ sera on normal spleen mononuclear cells (SMCs) and DQ transfected murine cell clones [Manuscript submitted for publication].

These 15 validated eplets are highlighted in the lightgreen rows. It appears that 9 of the 15 validated eplets have an HLA-EpiCheck score greater than 0.5, which represents a remarkable performance of HLA-EpiCheck. Out of these 9 eplets, 2 eplets have a score of 1, 4 eplets have a score in [0.8, 1[, and 3 eplets have a score in] 0.5, 0.8[. Regarding the remaining 6 out of the 15 evaluated eplets, 2 eplets have a score in [0.3, 0.5], 3 eplets have a score in] 0, 0.3[, and 1 eplet has a score of 0.

The 6 eplets with low prediction scores were further investigated. To do this, we checked the HLA-EpiCheck predictions of solvent-accessible neighbouring residues in the space of each eplet. We found that in the case of eplet 23L, residues 22, 25, 43 and 80 were predicted as epitope in the two antigens that display the eplet (see Table S8). In the case of eplet 125G, residue 126 was predicted to be an epitope in all 6 antigens displaying the eplet (see Table S9). These results suggest that the epitope is not centred on the eplet residues but in their vicinity. However, for the other 4 low scoring eplets, we did not find any neighbouring residues that were favourable for being epitope.

Eplet 75I is composed of residues 75I, 161D and 163I (74I, 160D and 162I in alleles DQA1*02, DQA1*04, DQA1*05 and DQA1*06). It is the only eplet with two completely different candidate epitope zones. This is because position 75/74 and positions 161/160 - 163/162 are approximately 50Å apart. Looking in detail at the >HLA-EpiCheck predictions for this eplet (see Table S10), we found that the position 75/74 was systematically predicted as epitope on all the antigens displaying the eplet. These results suggest that the patch around position 75/74 is the epitope actually recognized by antibodies.

## 4 Discussion

HLA-Epicheck uses an unprecedented dataset of short MD simulations to train a machine learning model that predicts B-cell epitopes on HLA proteins. The MD simulation data allows the integration of dynamic descriptors into the epitope prediction task, something that has never been done before. HLA-EpiCheck performs remarkably well in predicting epitopes at all HLA loci (although it performs less well at the locus DQ) and outperforms DiscoTope-3.0 on all metrics considered. Furthermore, when comparing HLA-EpiCheck predictions with the experimental evaluation of 15 non-confirmed eplets at the locus DQ, a remarkable consistency was found (9 out of 15 eplets validated experimentally were predicted positively by HLA-EpiCheck).

We split our descriptors into static and dynamic types to evaluate the contribution of each one to the performance of the tool. MDI values were used to perform a feature importance analysis. We found that hydrophobicity (static descriptor) plays a major role in the prediction capacity of HLA-EpiCheck. This result is in agreement with other studies that have pointed out the importance of hydrophobicity in antigen-antibody recognition [30–33]. Concerning the dynamic descriptors, the side-chain flexibility of the patch is ranked second in the feature importance analysis. This is consistent with what was reported in [17] where it is suggested that side-chain flexibility is key in antigen-antibody recognition. Additionally, the fact that the contribution of dynamic descriptors is about half of the prediction capacity of the tool supports the interest in using MD data for the epitope prediction task. This type of data, however, has the disadvantage of requiring heavy calculations for its generation.

The HLA-EpiCheck tool has some limitations that are worth bearing in mind. Our ML dataset presents several imbalances. One is the number of samples per label, as samples with the non-epitope label represent 69% of the dataset. This imbalance would explain the higher performance of the tool in predicting the non-epitope label. Another imbalance corresponds to the number of samples per locus, where the loci DR and C are the least represented in the dataset (495 and 536 samples respectively), while the locus B is by far the most represented (3351 samples). Interestingly, despite this imbalance, the performance of the tool at the loci DR and C is remarkably high (although with room for improvement especially at the locus DR). This is maybe related to the lower level of variability in DR and C as both loci only carry one polymorphic chain. Finally, we have the particular case of the locus DQ, where there is an imbalance in favor of the epitope label (486 samples for the epitope label and 107 samples for the non-epitope label). This imbalance would explain the drop in the ability of our tool to predict the non-epitope label of the locus DQ. According to the aforementioned limitations, efforts to improve HLA-EpiCheck should focus on the loci DQ and DR. For instance, we intend to expand our training set with new patches derived from DQ and DR antigens not considered so far.

Due to the major role of DQ antigens in the development of *de novo* DSA and thanks to an ongoing work aimed at experimental confirmation of DQ non-confirmed eplets, we were able to compare the results of HLA-EpiCheck predictions on a subset of non-confirmed eplets. The results show a notable consistency between the predictions of our tool and the experimental results. Additionally, we further investigated the 6 experimentally validated eplets that were not correctly predicted by our tool and found that for eplets 23L and 125G, HLA-EpiCheck predicts that the epitope could be centred on a residue in the vicinity of the eplet. Nevertheless, for the other 4 eplets, no similar explanation was found.Thus, these low scoring eplets can be considered as false negative with respect to HA-EpiCheck prediction. This is in agreement with our performance metrics that showed a precision greater than recall, suggesting that our tool can still be improved to reduce the amount of false-negative predicted samples. Alternatively, another hypothesis could be that the epitope induced by the eplet is located distantly from the eplet, in a region that we did not explore yet. Thus, one perspective for this work would be to perform a differential analysis between antigens differing only for these eplets in order to locate new candidate epitopes that are not centered or close to the eplet residues.

Regarding the case of eplet 75I, we found that the distance between position 75/74 and positions 161/160-163/162 indicates two completely different candidate epitope zones. However, the *𝒮*_*e-*_ *resid* scores suggest that the patch around position 75/74 is the true epitope.

Since confirmed eplet mismatches have been demonstrated to correlate with DSA formation and graft survival [24, 25], HLA-EpiCheck could be used for confirming non-confirmed eplets in the HLA Eplet Registry in order to minimize confirmed eplet mismatches when assigning transplants. Additionally, HLA-Epicheck could be used to perform a pairwise differential screening to identify new epitopes not yet described, in other words, to perform a differential comparison of predictions on donor and recipient HLA antigen pairs to identify new putative epitopes. Finally, adding new HLA antigens to the dataset beyond the 207 here simulated, is a perspective to enrich and potentially improve the performance of HLA-EpiCheck.

## 5 Conclusion

We introduce HLA-EpiCheck, a B-cell epitope predictor on HLA antigens. Short MD simulations of 207 HLA proteins were used to generate a dataset to train an Extra Trees binary classifier, i.e., a classifier to predict whether the patch associated with a solvent-accessible residue is an epitope or not. This dataset is composed of nearly 7000 patches (2117 epitopes and 4769 non epitopes) characterized by 18 descriptors of two types : static or dynamic. We performed a feature importance analysis to evaluate the contribution of each type to the performance of the tool and found that both groups of descriptors contribute almost equally. We also found that hydrophobicity and side-chain flexibility both play a key role in our predictor. The various evaluations presented in this paper suggest that our tool has a robust performance and that it has managed to capture patterns in the data leading to excellent results in all HLA sub-families. However, our tool still has room for improvement, especially with regard to reducing false-negative predictions. Furthermore, we compared HLA-EpiCheck against DiscoTope-3.0 (a state-of-the-art tool for B-cell epitope prediction) and found that our tool largely outperforms DiscoTope-3.0 on all metrics and HLA antigen sub-families. Finally, we compared the predictions of HLA-EpiCheck against experimental results obtained for 15 non-confirmed eplets of DQ antigens and found a remarkable performance of our tool. These results are particularly encouraging from the perspective of using HLA-EpiCheck so that non-confirmed eplets can be taken into account during donor-recipient HLA matching when allocating transplants.

## 6 Materials and Methods

### 6.1 Molecular Dynamics data generation

We selected 207 HLA antigens for molecular modeling, which include the antigens tested in Luminex Single Antigen (LSA) flow bead assays (see Table S1). Two procedures were implemented to generate the initial structures of the HLA antigens to run the MD simulations. One procedure corresponds to the antigens for which a structure exists in the PDB and the second one corresponds to antigens lacking a PDB structure (see Figure 3).

**Figure 3:**
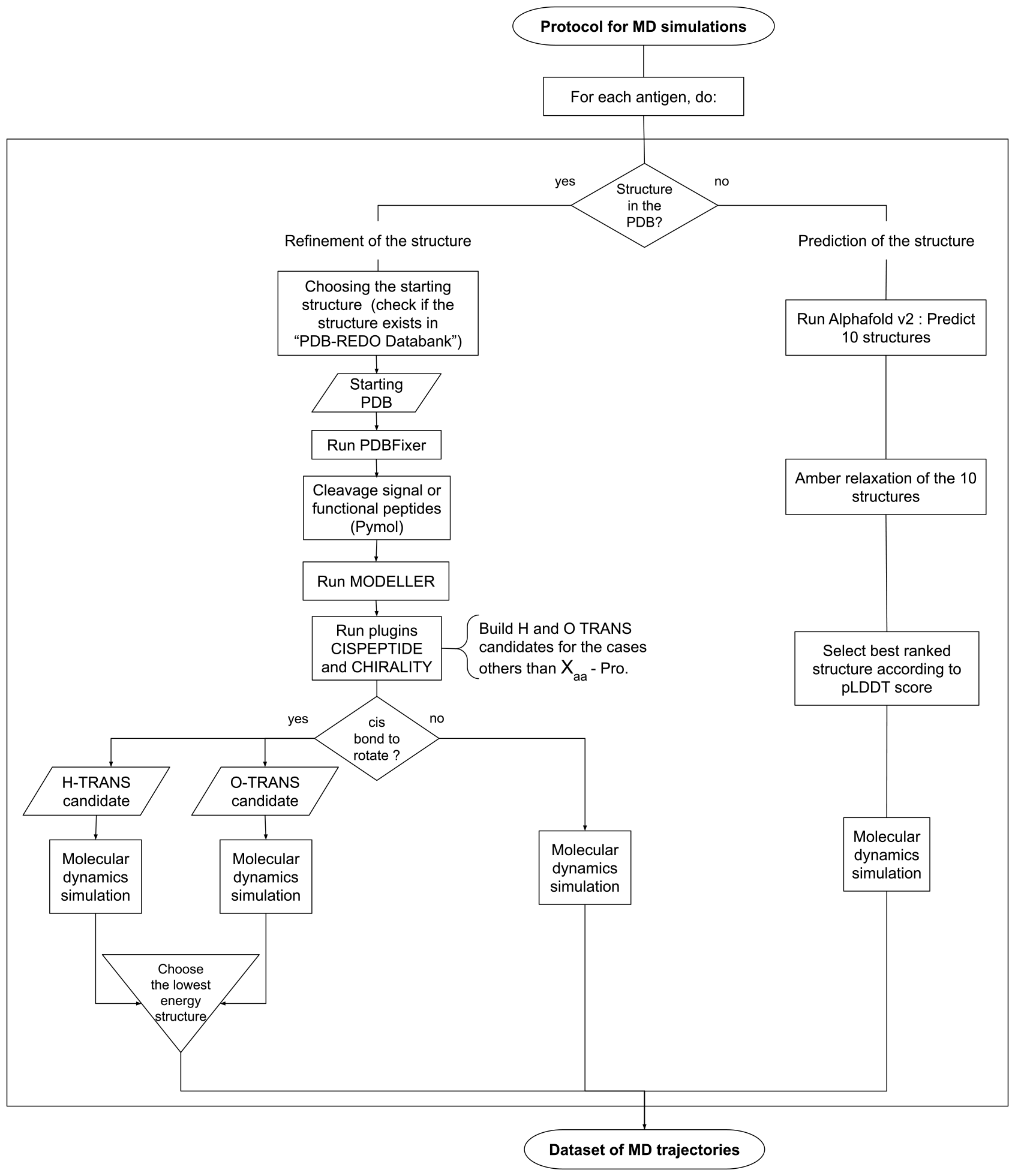
Protocol for MD simulations. There are two procedures, depending on whether the initial structure used to run the MD simulation comes from the PDB or from Alphafold.

#### 6.1.1 Identification of crystallographic structures in the PDB

The ≪ ncbiblast ≫ API [34] was used to find structures in the PDB using the HLA sequences in the Immuno Polymorphism Database (also known as IPD-IMGT/HLA database) [35]. Then, we checked if the structure was available in PDB REDO [36], otherwise, we downloaded the structure from the PDB (collected on June 23rd, 2020). The resolution values of the structures were obtained from the pdbe/EBI API [37] to finally choose a single template per antigen according to the best resolution.

#### 6.1.2 Refinement of previously identified PDB structures

PDBFixer [38] was used to identify and correct missing atoms or amino acids, remove small molecules and groove peptides. Pymol [39] was then used to remove signal peptides (if present) and trim the protein to keep only the extracellular globular part (see Table S2 for details). Loops and side chains were refined with MODELLER [40]. The Visual Molecular Dynamics (VMD) plugins CHIRALITY and CISPEPTIDE [41] were then used to check for chirality and cis-peptide bond errors. In cases where cis peptide bond errors not involving prolines were identified, structures were generated for hydrogen and oxygen rotations, then MD simulations were performed for both cases and an additional minimization stage was run to choose the conformation that minimized the energy of the system. Otherwise, a single MD simulation was run on the structure (see Figure 3).

#### 6.1.3 Generation of structures for antigens lacking any structure in the PDB

A local instance of Alphafold v2.1.0 was used to generate the structures of antigens absent from the PDB. Default parameters were utilized. Ten structures were generated per antigen to which an Amber relaxation was applied (see [18] for details) and a single structure was chosen based on the best confidence score (pLDDT).

#### 6.1.4 Molecular Dynamics simulations

MD simulation was performed for each antigen using the VMD [42] and NAMD3 tools. For the generation of the system topology, the VMD AutoPSF tool was used together with the CHARMM36 force field [43]. The system was solvated using the TIP3P explicit solvent model and then neutralized using the VMD AutoIonize tool. The Particle Mesh Ewald (PME) method was used to calculate the electrostatic energy with a distance truncation of 11 Å. The simulation was carried out under NPT conditions, that is, constant pressure (1 atm) and temperature (310 K) and was composed of 2 stages: (i) minimization, in which the system was brought into room temperature (310K), (ii) system equilibration of ∼10ns. 500 frames from the last 5ns of simulations (also known as trajectories) were used for subsequent analyses. It is worth mentioning that short MD simulations were performed because as demonstrated elsewhere [17], side-chain motions are fast enough to be studied through short simulations. See Supplementary Materials for data availability.

### 6.2 Machine Learning dataset generation

To build our ML dataset, we defined the epitopes by patches centered on a solvent-accessible residue. Then a set of 18 descriptors/features (see Table 4) are computed for each patch and used for ML.

**Table 4:**
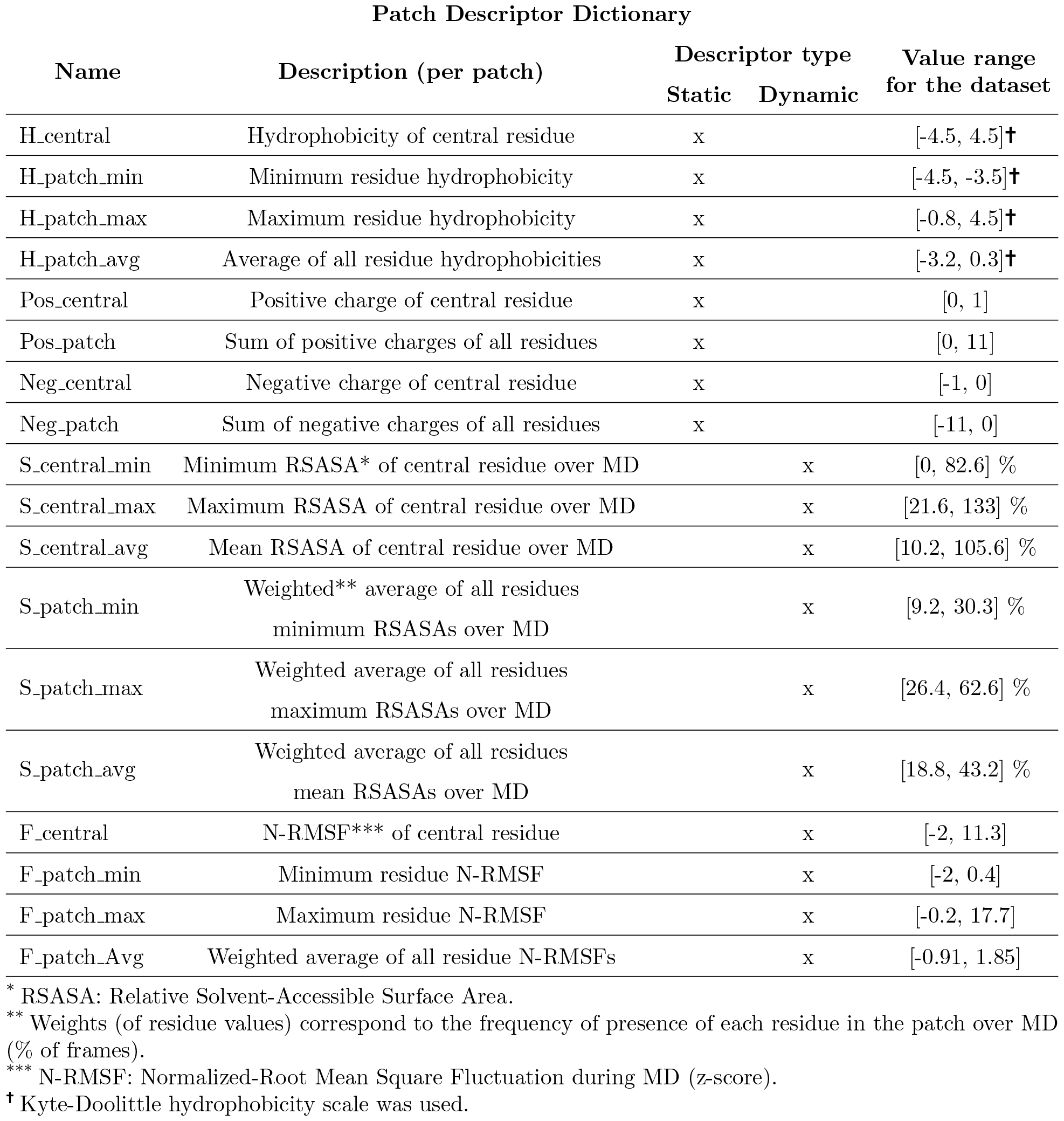
Descriptor dictionary of the ML dataset.

#### 6.2.1 Definition of epitope and non-epitope patches

The 3D objects considered in our ML approach are fragments of the protein surface called ≪ patches ≫. In this study, a patch is centered on a solvent-accessible residue and contains all other solvent-accessible residues present in a radius of 15Å from the central residue (distances are computed between the centers of mass of the residues). We used the average Relative Solvent Accessible Surface Area (RSASA) along the trajectory to define whether a residue is solvent accessible. Namely, a residue is considered solvent accessible when its average RSASA along the trajectory is higher than 20%. SASA calculator implemented in VMD [44] was utilized to compute the absolute solvent accessibility of residues and the theoretical ≪ ALL ≫ maximum solvent accessibility of each residue defined in [45] (see Table S3) was used to calculate RSASA values for each frame.

From the HLA Eplet Registry, we obtained a list of antibody-verified eplets. Epitope patches were computed as centered on solvent-accessible residues from this list (here termed as eplet residues).

The non-epitope patches were centered on a solvent-accessible residue that had not been listed as part of an eplet in the HLA Eplet Registry (either antibody-verified or not). However, not all such patches were eligible as negative counterparts of epitope patches. A first condition was that, in addition to the central residue, none of the solvent-accessible residues composing a non-epitope patch should be part of an eplet (either antibody verified or not). Another condition was that none of the solvent-accessible residues composing a non-epitope patch should be in a locus-conserved position. Indeed, considering locus-conserved positions could be misleading as these positions could by chance display B-cell epitope behavior but cannot be recognized as such because they cannot raise immune response as they are conserved throughout all individuals. Thus, considering these locus-conserved positions could bias the training by introducing ≪ silencious ≫ positive examples among the non-epitope ones.

Homemade Pymol and Python scripts were written to compute the patches (see supplementary material).

#### 6.2.2 Definition and calculation of patch descriptors

##### 2.2.2.1 Static descriptors

Static descriptors are attached to a patch and do not vary along MD simulation. We computed residue hydrophobicity according to the Kyte-Doolittle scale [46] and derived 4 descriptors for a patch: the hydrophobicity of the central residue, the minimum, maximum and average residue hydrophobicity over all the residues of the patch.

The other static descriptors were derived from the residue electrostatic charges namely a negative charge for aspartate (D) and glutamate (E) residues, and a positive charge for lysine (K) and arginine (R) residues. Four descriptors were computed for each patch, two for the negative charges and two for the positive charges. For each type of charge (either negative or positive), we computed (i) the charge of the central residue (the attribute Pos central takes the value +1 for a positively charged residue, otherwise 0, and the same for the attribute Neg central except that the value is -1 for a negatively charged residue), (ii) the sum of the charges (either positive or negative) of the residues in the patch (integer ≥ 0 for the attribute Pos patch,, integer ≤ 0 for the attribute Neg patch).

#### 2.2.2.2 Dynamic descriptors

Dynamic variables measure conformational changes of the patch structure during MD simulation. A first group of 6 descriptors reflected the variation of the RSASA percentage during the run. RSASA values were calculated per residue for each of the 500 frames considered. For the central residue of a patch, we kept the minimum, maximum and mean RSASA values along the run. For the entire patch, we used the minimum, maximum and mean RSASA values of each residue in the patch along the run and we aggregated these values according to the type (all minima, all maxima, all means) as three weighted averages in which the weights corresponded for each residue to the % of frames in which this residue was really present in the patch (distance between centers of mass ≤ 15Å).

A second group of 4 descriptors reflected side-chain flexibility along the run. These descriptors were derived from the Normalised Root Mean Squared Fluctuation (N-RMSF) values [17]. One corresponded to the N-RMSF of the central residue of a patch and the others to the minimum, maximum and weighted average N-RMSF over all residues of the patch (weights are the same as defined above). RMSF values were calculated from the 500 frames considered for each trajectory using VMD. Z-score normalisation method was used where only solvent-accessible residues were included and five residues at each terminus were discarded because of their artefactual flexibility.

### 6.2.3 Dataset composition and evaluation metrics

HLA-EpiCheck is a binary classifier that predicts whether the patch centered on a solvent-accessible residue is an epitope or not. It is important to highlight that the dataset generated here presents an important imbalance between labels (non-epitope/epitope ratio ∼0.44) as can be seen in Table 5. Therefore, the precision and recall metrics are more relevant to evaluate the ability of a binary classifier to predict a label. We used the F1 score (harmonic mean of the precision and recall), Area Under the Curve of the Receiver Operating Characteristic curve (AUC-ROC), Area Under the Curve of the Precision-Recall curve (AUC-PR) and Matthews Correlation Coefficient (MCC) metrics to evaluate our predictor [47]. Scikit-Learn [48] was used to compute all the metrics.

**Table 5:** Overview of the ML dataset. HLA antigens are divided in A, B, C, DP, DQ and DR sub-families. These sub-families come from the loci encoding these proteins.

|  |  | A | B | C | DP | DQ | DR | Total |
| --- | --- | --- | --- | --- | --- | --- | --- | --- |
| Dataset | # antigens | 34 | 59 | 17 | 31 | 30 | 36 | **207** |
|  | # epitope labels | 434 | 529 | 177 | 184 | 486 | 307 | **2117** |
|  | # non-epitope labels | 656 | 2822 | 359 | 637 | 107 | 188 | **4769** |
| Training set (80%) | # antigens | 27 | 47 | 13 | 24 | 24 | 28 | **163** |
|  | # epitope labels | 353 | 427 | 134 | 138 | 395 | 241 | **1688** |
|  | # non-epitope labels | 506 | 2270 | 281 | 529 | 95 | 141 | **3822** |
| Test set (20%) | # antigens | 7 | 12 | 4 | 7 | 6 | 8 | **44** |
|  | # epitope labels | 81 | 102 | 43 | 46 | 91 | 66 | **429** |
|  | # non-epitope labels | 150 | 552 | 78 | 108 | 12 | 47 | **947** |

### 6.3 Training of the epitope predictor tool

The Scikit-Learn implementation of the Extra Trees [49] method was employed to train HLA-EpiCheck. The Extra Trees method is an ensemble learning method. It builds multiple decision trees by selecting random subsets of features but instead of looking for the most discriminative threshold, it is randomly generated for each candidate feature and the best of these randomly-generated thresholds is picked as the splitting rule. In contrast to the original publication [49], the Scikit-Learn implementation combines trees by averaging their probabilistic predictions, instead of letting each tree vote for a single label. Thus, the final predicted label is the one with the highest probability. Other learning algorithms were tested (results not shown) such as logistic regression, support vector machines, gradient boosting classifiers and multi-layer perceptrons but Extra Trees was the best-performing of all. For training the predictor, 80% of the dataset (training set) was used, while for the evaluation of the performance (test set) the remaining 20% was used. The hyperparameters of the predictor were defined with the help of the gridsearchCV function of Scikit-Learn, for which the training set, a 5-folds cross-validation and the F1 metric were utilized. Table S4 shows the parameters finally kept for training the predictor. For the performance evaluation on the training set, 10 repetitions of 10-fold cross-validations were calculated. For the feature importance analysis, a built-in function in the ExtraTreesClassifier class of Scikit-Learn was used to compute Mean Decrease in Impurity (MDI) [50] values.

### 6.4 HLA-EpiCheck score at eplet level for comparison with experimental results

Since HLA-EpiCheck acts on patches centered on a unique residue while some eplets are composed of more than one solvent-accessible residue, and because the prediction can be performed on all HLA antigens displaying an eplet of interest, we had to aggregate the results obtained by patch prediction for each eplet residue and for all antigens displaying this eplet. This led to the definition of an HLA-EpiCheck score. To define this score, we first inspected the distance between residues in cases of eplets composed of more than one solvent-accessible residue, hereafter named ≪ composite ≫ eplets. We did this inspection on all non-confirmed eplets of the locus DQ and found that in all eplets except one (75I), the eplet residues are less than 9Å apart. For this reason, we defined two scores. The first, named *𝒮*_*e*_, will aggregate predictions over all the eplet residues as well as predictions over all the antigens displaying the eplet, i.e., the score will be per eplet. The second score, named *𝒮*_*e_resid*_, will aggregate only the predictions over all the antigens displaying an eplet residue, i.e., the score will be per eplet residue. Having said that, we defined the scores as follows:

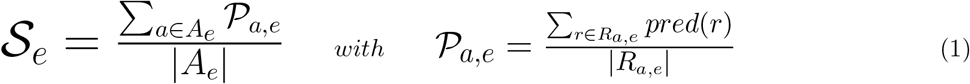

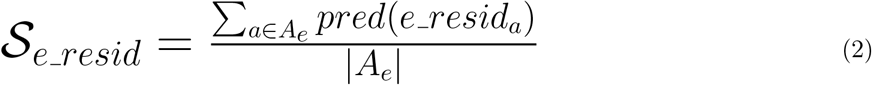

where *𝒮*_*e*_ is the score for the eplet *e, A*_*e*_ is the set of antigens carrying the eplet *e, P*_*a,e*_ is the prediction score aggregated on all solvent-accessible residues members of eplet *e* for antigen *a, R*_*a,e*_ is the set of solvent-accessible residues members of eplet *e* on antigen *a, pred*(*r*) is the HLA-EpiCheck prediction on residue *r* (0 for a non-epitope prediction and 1 for an epitope prediction), *𝒮*_*e_resid*_ is the score for the eplet residue *e_resid* of the eplet *e* and *pred*(*e_resid*_*a*_) is the HLA-EpiCheck prediction for the eplet residue *e_resid* on antigen *a*. These scores range from 0 to 1 and we will consider that a non-confirmed eplet or eplet residue is favorable to antibody recognition if the corresponding score is greater than 0.5.

## Supporting information

All supplemental data

## 7 Acknowledgments

Experiments presented in this paper were carried out using the Grid’5000 testbed, supported by a scientific interest group hosted by Inria and including CNRS, RENATER and several Universities as well as other organizations (see https://www.grid5000.fr), as well as the MBI-DS4H platform hosted by Inria/Loria and funded by CPER IT2MP (Contrat Plan État Région, Innovations, Technologiques, Modélisation & Médicine Personalisée), including a FEDER co-funding.

## Notes

### Competing Interest Statement

The authors have declared no competing interest.

