## Supplementary material for "HLA-EpiCheck : A B-cell epitope prediction tool for HLA proteins using molecular dynamics simulation data": All supplemental data

### 8 Supplementary material

**Table S1:** List of modeled HLA antigens. DP and DQ antigens are composed of two polymorphic protein chains, produced from the DPA1 and DPB1, or DQA1 and DQB1 loci, respectively.

| Locus | Antigens |
|---|---|
| A | A*01:01, A*02:01, A*02:02, A*02:03, A*02:05, A*02:06, A*02:07, A*03:01, A*11:01, A*11:02, A*23:01, A*24:02, A*24:03, A*25:01, A*26:01, A*29:01, A*29:02, A*30:01, A*30:02, A*30:03, A*31:01, A*32:01, A*33:01, A*33:03, A*34:01, A*36:01, A*43:01, A*66:01, A*66:02, A*68:01, A*68:02, A*69:01, A*74:01, A*80:01 |
| B | B*07:02, B*08:01, B*13:01, B*13:02, B*14:01, B*14:02, B*15:01, B*15:02, B*15:03, B*15:10, B*15:11, B*15:12, B*15:13, B*15:16, B*18:01, B*27:03, B*27:04, B*27:05, B*27:06, B*27:08, B*27:09, B*35:01, B*35:08, B*37:01, B*38:01, B*39:01, B*40:01, B*40:02, B*40:06, B*41:01, B*41:03, B*41:04, B*42:01, B*42:02, B*44:02, B*44:03, B*44:05, B*45:01, B*46:01, B*47:01, B*48:01, B*49:01, B*50:01, B*51:01, B*51:02, B*52:01, B*53:01, B*54:01, B*55:01, B*56:01, B*57:01, B*57:03, B*58:01, B*59:01, B*67:01, B*73:01, B*78:01, B*81:01, B*82:01 |
| C | C*01:02, C*02:02, C*03:02, C*03:03, C*03:04, C*04:01, C*05:01, C*06:02, C*07:02, C*08:01, C*08:02, C*12:03, C*14:02, C*15:02, C*16:01, C*17:01, C*18:02 |
| DP | DPA1*01:03-DPB1*01:01, DPA1*01:03-DPB1*02:01, DPA1*01:03-DPB1*03:01, DPA1*01:03-DPB1*04:01, DPA1*01:03-DPB1*04:02, DPA1*01:03-DPB1*06:01, DPA1*01:03-DPB1*11:01, DPA1*01:03-DPB1*19:01, DPA1*01:03-DPB1*23:01, DPA1*01:03-DPB1*28:01, DPA1*01:04-DPB1*18:01, DPA1*01:05-DPB1*03:01, DPA1*01:05-DPB1*18:01, DPA1*01:05-DPB1*28:01, DPA1*02:01-DPB1*01:01, DPA1*02:01-DPB1*03:01, DPA1*02:01-DPB1*05:01, DPA1*02:01-DPB1*06:01, DPA1*02:01-DPB1*09:01, DPA1*02:01-DPB1*13:01, DPA1*02:01-DPB1*14:01, DPA1*02:01-DPB1*15:01, DPA1*02:01-DPB1*17:01, DPA1*02:01-DPB1*18:01, DPA1*02:02-DPB1*05:01, DPA1*02:02-DPB1*10:01, DPA1*02:02-DPB1*11:01, DPA1*02:02-DPB1*13:01, DPA1*03:01-DPB1*13:01, DPA1*03:01-DPB1*20:01, DPA1*04:01-DPB1*28:01 |
| DQ | DQA1*01:01-DQB1*05:01, DQA1*01:01-DQB1*06:02, DQA1*01:02-DQB1*05:01, DQA1*01:02-DQB1*05:02, DQA1*01:02-DQB1*06:02, DQA1*01:02-DQB1*06:04, DQA1*01:02-DQB1*06:09, DQA1*01:03-DQB1*06:01, DQA1*01:03-DQB1*06:03, DQA1*02:01-DQB1*02:01, DQA1*02:01-DQB1*02:02, DQA1*02:01-DQB1*03:01, DQA1*02:01-DQB1*03:02, DQA1*02:01-DQB1*03:03, DQA1*02:01-DQB1*04:01, DQA1*02:01-DQB1*04:02, DQA1*03:01-DQB1*02:01, DQA1*03:01-DQB1*03:01, DQA1*03:01-DQB1*03:02, DQA1*03:01-DQB1*03:03, DQA1*03:02-DQB1*03:02, DQA1*03:02-DQB1*03:03, DQA1*03:03-DQB1*04:01, DQA1*04:01-DQB1*02:01, DQA1*04:01-DQB1*04:02, DQA1*05:01-DQB1*02:01, DQA1*05:03-DQB1*03:01, DQA1*05:05-DQB1*03:01, DQA1*05:08-DQB1*02:01, DQA1*06:01-DQB1*03:01 |
| DR | DRB1*01:01, DRB1*01:02, DRB1*01:03, DRB1*03:01, DRB1*03:02, DRB1*04:01, DRB1*04:02, DRB1*04:03, DRB1*04:04, DRB1*04:05, DRB1*07:01, DRB1*08:01, DRB1*09:01, DRB1*09:02, DRB1*10:01, DRB1*11:01, DRB1*11:04, DRB1*12:01, DRB1*12:02, DRB1*13:01, DRB1*13:03, DRB1*14:01, DRB1*14:02, DRB1*14:54, DRB1*15:01, DRB1*15:02, DRB1*15:03, DRB1*16:01, DRB1*16:02, DRB3*01:01, DRB3*02:02, DRB3*03:01, DRB4*01:01, DRB4*01:03, DRB5*01:01, DRB5*02:02 |

**Table S2:** Length of modeled sequences per locus. Only the extracellular globular part of the protein was modeled, so the signal peptide, the amino acids linking the extracellular globular part to the transmembrane part, the transmembrane part and the intracellular part were excluded.

| **locus** | **A, B, C** | | **DP** | | **DQ** | | **DR** | |
|---|---|---|---|---|---|---|---|---|
| **chain** | **chain $\alpha$** | **$\beta$2-microglobulin** | **chain $\alpha$** | **chain $\beta$** | **chain $\alpha$** | **chain $\beta$** | **chain $\alpha$** | **chain $\beta$** |
| **modeled length** | 276 | 99 | 183 | 191 | 186 | 192 | 182 | 192 |

**Table S3:** Maximum solvent accessibility of residues [45]

| **Residue** | **Max. accessibility** [Å$^2$] |
|---|---|
| ALA | 138 |
| ARG | 285 |
| ASN | 204 |
| ASP | 204 |
| CYS | 169 |
| GLU | 233 |
| GLN | 234 |
| GLY | 114 |
| HIS | 231 |
| ILE | 208 |
| LEU | 211 |
| LYS | 246 |
| MET | 227 |
| PHE | 251 |
| PRO | 166 |
| SER | 161 |
| THR | 182 |
| TRP | 295 |
| TYR | 274 |
| VAL | 184 |

**Table S4:** Hyperparameters of the Extra Trees model

| **Hyperparameter** | **Value** |
|---|---|
| Criterion | gini |
| n_estimators (trees) | 100 |
| min_samples_leaf | 1 |
| min_samples_split | 2 |

**Table S5:** Performance evaluation on the training set carrying out 10 repetitions of 10-fold cross-validations. Precision, Recall and F1 values were computed for each repetition as well as global values across all repetitions.

| **Repetition** | **Precision** | | | | **Recall** | | | | **F1** | | | |
|---|---|---|---|---|---|---|---|---|---|---|---|---|
| | **Epitope** | | **Non-epitope** | | **Epitope** | | **Non-epitope** | | **Epitope** | | **Non-epitope** | |
| | **mean** | **std.** | **mean** | **std.** | **mean** | **std.** | **mean** | **std.** | **mean** | **std.** | **mean** | **std.** |
| 0 | 0.922 | 0.015 | 0.934 | 0.011 | 0.842 | 0.026 | 0.967 | 0.007 | 0.879 | 0.016 | 0.951 | 0.007 |
| 1 | 0.916 | 0.035 | 0.935 | 0.008 | 0.836 | 0.034 | 0.966 | 0.010 | 0.873 | 0.012 | 0.950 | 0.005 |
| 2 | 0.922 | 0.013 | 0.934 | 0.013 | 0.833 | 0.028 | 0.969 | 0.013 | 0.875 | 0.016 | 0.951 | 0.009 |
| 3 | 0.914 | 0.026 | 0.935 | 0.011 | 0.838 | 0.026 | 0.968 | 0.010 | 0.874 | 0.023 | 0.951 | 0.008 |
| 4 | 0.920 | 0.024 | 0.933 | 0.008 | 0.843 | 0.029 | 0.966 | 0.010 | 0.880 | 0.021 | 0.949 | 0.007 |
| 5 | 0.925 | 0.030 | 0.934 | 0.007 | 0.837 | 0.028 | 0.967 | 0.007 | 0.878 | 0.017 | 0.950 | 0.006 |
| 6 | 0.919 | 0.024 | 0.930 | 0.016 | 0.832 | 0.034 | 0.969 | 0.010 | 0.873 | 0.024 | 0.949 | 0.010 |
| 7 | 0.920 | 0.023 | 0.934 | 0.018 | 0.840 | 0.021 | 0.968 | 0.012 | 0.878 | 0.018 | 0.951 | 0.008 |
| 8 | 0.924 | 0.020 | 0.935 | 0.017 | 0.834 | 0.022 | 0.965 | 0.010 | 0.877 | 0.018 | 0.950 | 0.011 |
| 9 | 0.916 | 0.026 | 0.938 | 0.015 | 0.839 | 0.020 | 0.966 | 0.008 | 0.876 | 0.021 | 0.952 | 0.008 |
| **Global** | **0.920** | **0.025** | **0.934** | **0.013** | **0.837** | **0.028** | **0.967** | **0.010** | **0.876** | **0.019** | **0.950** | **0.008** |

**Table S6:** Performance evaluation on the training set carrying out 10 repetitions of 10-fold cross-validations. AUC-ROC, AUC-PR and MCC values were computed for each repetition as well as global values across all repetitions.

| **Repetition** | **AUC-ROC** | | **AUC-PR** | | **MCC** | |
|---|---|---|---|---|---|---|
| | **mean** | **std.** | **mean** | **std.** | **mean** | **std.** |
| 0 | 0.9801 | 0.0064 | 0.9575 | 0.0081 | 0.8301 | 0.0147 |
| 1 | 0.9785 | 0.0064 | 0.9589 | 0.0092 | 0.8348 | 0.0323 |
| 2 | 0.9796 | 0.0037 | 0.9580 | 0.0097 | 0.8283 | 0.0240 |
| 3 | 0.9796 | 0.0040 | 0.9592 | 0.0134 | 0.8313 | 0.0215 |
| 4 | 0.9796 | 0.0046 | 0.9593 | 0.0070 | 0.8341 | 0.0191 |
| 5 | 0.9784 | 0.0061 | 0.9605 | 0.0120 | 0.8291 | 0.0268 |
| 6 | 0.9805 | 0.0058 | 0.9578 | 0.0069 | 0.8381 | 0.0271 |
| 7 | 0.9796 | 0.0052 | 0.9581 | 0.0099 | 0.8239 | 0.0260 |
| 8 | 0.9782 | 0.0057 | 0.9582 | 0.0117 | 0.8225 | 0.0195 |
| 9 | 0.9791 | 0.0039 | 0.9580 | 0.0111 | 0.8273 | 0.0281 |
| **Global** | **0.9793** | **0.0053** | **0.9586** | **0.0101** | **0.8299** | **0.0251** |

**Table S7:** List of non-confirmed eplets for the locus DQ in HLA Eplet Registry (accessed February, 2023)

| No | Eplet name | Eplet description |
|---|---|---|
| 1 | 2G | 2G 199T |
| 2 | 3P | 3P 9L 37D |
| 3 | 3S | 3S |
| 4 | 9F | 9F |
| 5 | 9Y | 9Y |
| 6 | 13GM | 13G 14M |
| 7 | 23L | 23L |
| 8 | 23R | 23R |
| 9 | 25FT | 25F 26T |
| 10 | 25YT | 25Y 26T |
| 11 | 26L | 26L |
| 12 | 30H | 30H |
| 13 | 37YA | 37Y 38A |
| 14 | 37YV | 37Y 38V |
| 15 | 40ERV | 40E 41R 45V |
| 16 | 45G | 45G |
| 17 | 55PPA | 55P 56P 57A |
| 18 | 55PPD | 55P 56P 57D |
| 19 | 55RPD | 55R 56P 57D |
| 20 | 56P | 56P |
| 21 | 56PA | 56P 57A |
| 22 | 56PD | 56P 57D |
| 23 | 56PS | 56P 57S |
| 24 | 66D | 66D 67I |
| 25 | 66DR | 66D 67I 70R |
| 26 | 66ER | 66E 67V 70R 71T |
| 27 | 66EV | 66E 67V |
| 28 | 66IL | 66I 69L |
| 29 | 66IT | 66I 69T |
| 30 | 67VG | 66E 67V 70G |
| 31 | 67VT | 67V 71T |
| 32 | 70GT | 67V 70G 71T |
| 33 | 70RT | 70R 71T |
| 34 | 74EL | 74E 75L 77T |
| 35 | 75I | 75I1 61D 163I |
| 36 | 75IL | 75I 76L |
| 37 | 75V | 75V |
| 38 | 76L | 76L |
| 39 | 86A | 86A |
| 40 | 125G | 125G |
| 41 | 129H | 129H |
| 42 | 129QS | 129Q 130S |
| 43 | 130A | 130A |
| 44 | 130Q | 86G 130Q |
| 45 | 130R | 130R |
| 46 | 135D | 135D |
| 47 | 135G | 135G |
| 48 | 160A | 160A |
| 49 | 160AD | 160A 161D |
| 50 | 160D | 160D |
| 51 | 160S | 160S |
| 52 | 167H | 167H |
| 53 | 167R | 167R |
| 54 | 175E | 175E |
| 55 | 185I | 185I |
| 56 | 185T | 185T |

**Table S8:** HLA-EpiCheck predictions on the eplet 23L and its neighboring residues. Columns **Ag1** and **Ag2** correspond to predictions on antigens DQA1*02:01-DQB1*04:01 and DQA1*03:03-DQB1*04:01 respectively. Missing prediction value corresponds to a non solvent-accessible residue.

| **Central residue** | **Chain** | **Ag1** | **Ag2** |
|---|---|---|---|
| 23L | $\beta$ | 1 | 0 |
| 3I | $\alpha$ | - | 0 |
| 4V | $\alpha$ | 0 | 0 |
| 19N | $\beta$ | 1 | 0 |
| 22E | $\beta$ | 1 | 1 |
| 25R | $\beta$ | 1 | 1 |
| 43D | $\beta$ | 1 | 1 |
| 80R | $\beta$ | 1 | 1 |

**Table S9:** HLA-Epicheck predictions on the eplet 125G and its neighboring residues. Columns **Ag1**, **Ag2**, **Ag3**, **Ag4**, **Ag5** and **Ag6** correspond to predictions on antigens DQA1*01:01-DQB1*06:02, DQA1*01:02-DQB1*06:02, DQA1*01:02-DQB1*06:04, DQA1*01:02-DQB1*06:09, DQA1*01:03-DQB1*06:01 and DQA1*01:03-DQB1*06:03 respectively. Missing prediction values correspond to non solvent-accessible residues.

| **Central residue** | **Chain** | **Ag1** | **Ag2** | **Ag3** | **Ag4** | **Ag5** | **Ag6** |
|---|---|---|---|---|---|---|---|
| 125G | $\beta$ | 0 | - | - | - | - | - |
| 124P | $\beta$ | 0 | - | 1 | 1 | 1 | 1 |
| 126Q | $\beta$ | 1 | 1 | 1 | 1 | 1 | 1 |
| 147L | $\beta$ | 0 | 0 | 0 | 0 | 1 | 0 |

**Table S10:** HLA-Epicheck predictions before aggregation for all antigens displaying the eplet 75I. NA=Non-applicable.

| No | Antigen | 74I | 75I | 160D | 161D | 162I | 163I |
|---|---|---|---|---|---|---|---|
| 1 | DQA1*01:01-DQB1*05:01 | NA | 1 | NA | 0 | NA | 0 |
| 2 | DQA1*01:01-DQB1*06:02 | NA | 1 | NA | 0 | NA | 0 |
| 3 | DQA1*01:02-DQB1*05:01 | NA | 1 | NA | 1 | NA | 1 |
| 4 | DQA1*01:02-DQB1*05:02 | NA | 1 | NA | 0 | NA | 0 |
| 5 | DQA1*01:02-DQB1*06:02 | NA | 1 | NA | 1 | NA | 1 |
| 6 | DQA1*01:02-DQB1*06:04 | NA | 1 | NA | 1 | NA | 1 |
| 7 | DQA1*01:02-DQB1*06:09 | NA | 1 | NA | 0 | NA | 0 |
| 8 | DQA1*01:03-DQB1*06:01 | NA | 1 | NA | 1 | NA | 1 |
| 9 | DQA1*01:03-DQB1*06:03 | NA | 1 | NA | 1 | NA | 1 |
| 10 | DQA1*02:01-DQB1*02:01 | 1 | NA | 0 | NA | 1 | NA |
| 11 | DQA1*02:01-DQB1*02:02 | 1 | NA | 1 | NA | 1 | NA |
| 12 | DQA1*02:01-DQB1*03:01 | 1 | NA | 1 | NA | 1 | NA |
| 13 | DQA1*02:01-DQB1*03:02 | 1 | NA | 0 | NA | 0 | NA |
| 14 | DQA1*02:01-DQB1*03:03 | 1 | NA | 0 | NA | 0 | NA |
| 15 | DQA1*02:01-DQB1*04:01 | 1 | NA | 0 | NA | 0 | NA |
| 16 | DQA1*02:01-DQB1*04:02 | 1 | NA | 0 | NA | 0 | NA |
| 17 | DQA1*03:01-DQB1*02:01 | NA | 1 | NA | 0 | NA | 1 |
| 18 | DQA1*03:01-DQB1*03:01 | NA | 1 | NA | 0 | NA | 1 |
| 19 | DQA1*03:01-DQB1*03:02 | NA | 1 | NA | 1 | NA | 0 |
| 20 | DQA1*03:01-DQB1*03:03 | NA | 1 | NA | 0 | NA | 1 |
| 21 | DQA1*03:02-DQB1*03:02 | NA | 1 | NA | 0 | NA | 0 |
| 22 | DQA1*03:02-DQB1*03:03 | NA | 1 | NA | 1 | NA | 1 |
| 23 | DQA1*03:03-DQB1*04:01 | NA | 1 | NA | 0 | NA | 1 |
| 24 | DQA1*04:01-DQB1*02:01 | 1 | NA | 0 | NA | 1 | NA |
| 25 | DQA1*04:01-DQB1*04:02 | 1 | NA | 1 | NA | 1 | NA |
| 26 | DQA1*06:01-DQB1*03:01 | 1 | NA | 0 | NA | 1 | NA |

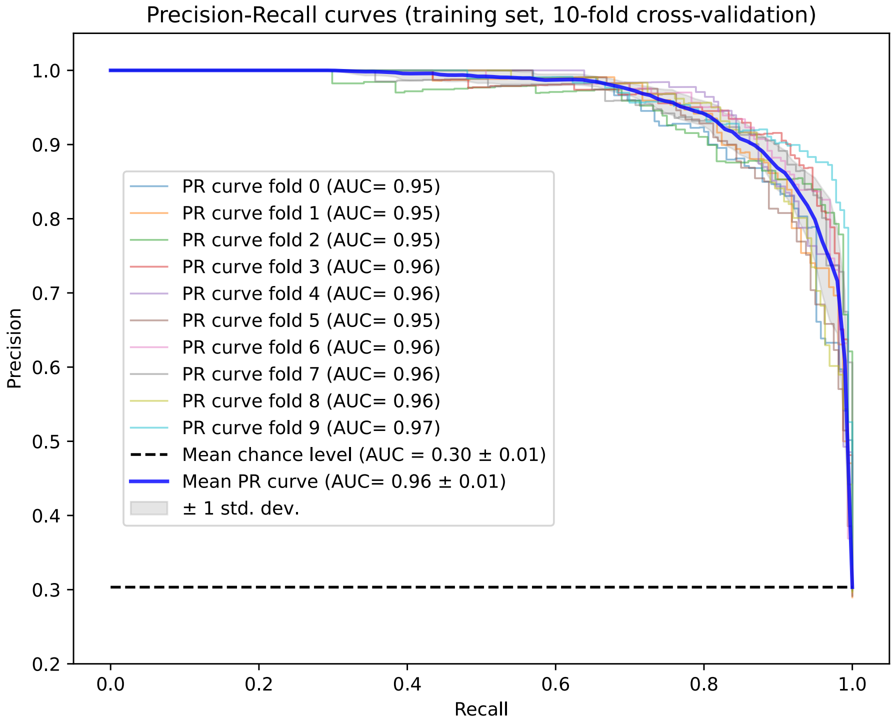


**Figure S1:** Prediction-Recall (PR) curves of a 10-fold cross-validation on training set (repetition 0). AUC value was computed for each fold as well as for the mean PR curve.

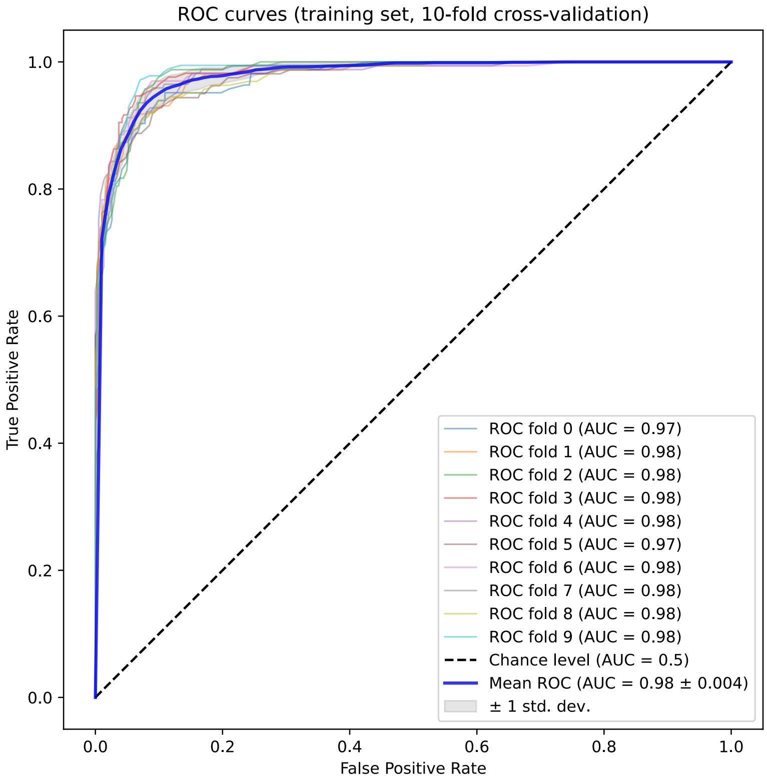


**Figure S2:** Receiver Operating Characteristic (ROC) curves of a 10-fold cross-validation on training set (repetition 0). AUC value was computed for each fold as well as for the mean ROC curve.

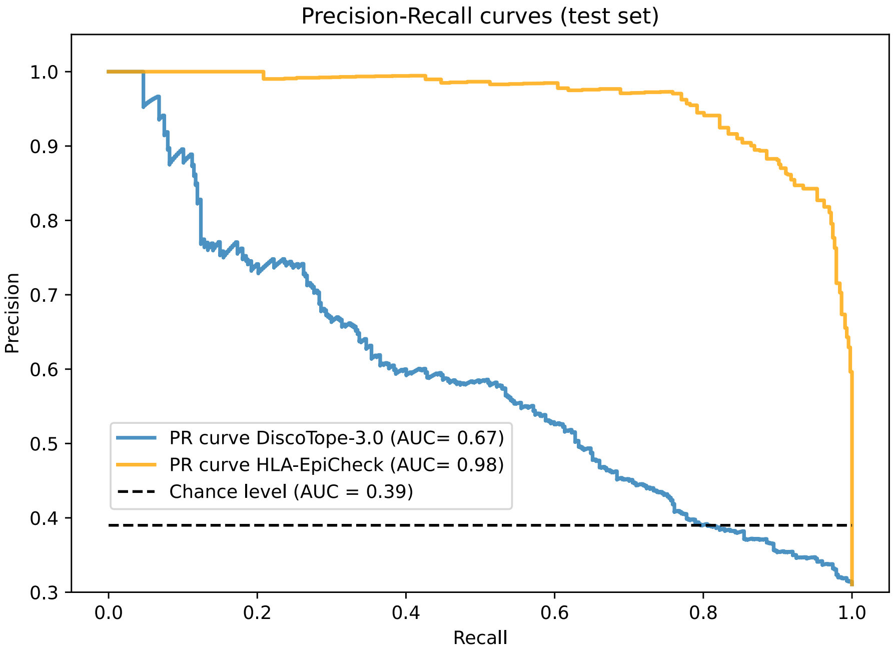


**Figure S3:** Prediction-Recall (PR) curves for HLA-EpiCheck and DiscoTope-3.0 on test set.

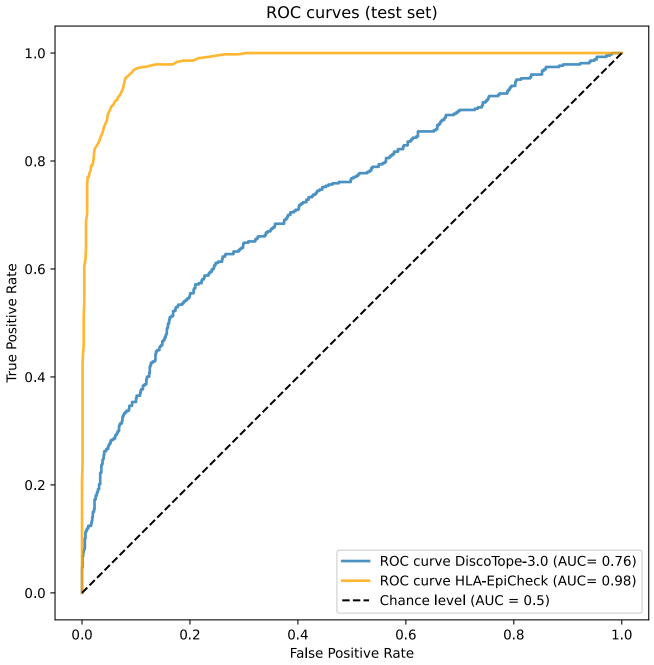


**Figure S4:** ROC curves for HLA-EpiCheck and DiscoTope-3.0 on test set.

[39] The PyMOL Molecular Graphics System [software]. Schrödinger;. Available from: http://www.pymol.org.

[40] Šali A, Blundell TL. Comparative Protein Modelling by Satisfaction of Spatial Restraints. Journal of Molecular Biology. 1993. Available from: https://doi.org/10.1006/jmbi.1993.1626.

[41] Schreiner E, Trabuco LG, Freddolino PL, Schulten K. Stereochemical errors and their implications for molecular dynamics simulations. BMC Bioinformatics. 2011. Available from: https://doi.org/10.1186/1471-2105-12-190.

[42] Humphrey W, Dalke A, Schulten K. VMD: Visual molecular dynamics. Journal of Molecular Graphics. 1996. Available from: https://doi.org/10.1016/0263-7855(96)00018-5.

[43] Lee J, Cheng X, Swails JM, Yeom MS, Eastman PK, Lemkul JA, et al. CHARMM-GUI Input Generator for NAMD, GROMACS, AMBER, OpenMM, and CHARMM/OpenMM Simulations Using the CHARMM36 Additive Force Field. Journal of Chemical Theory and Computation. 2016. Available from: https://doi.org/10.1021/acs.jctc.5b00935.

[44] Varshney A, Brooks FP, Wright WV. Linearly Scalable Computation of Smooth Molecular Surfaces. IEEE Computer Graphics and Applications. 1994.
